# Heme synthesis inhibition blocks angiogenesis via mitochondrial dysfunction

**DOI:** 10.1101/836304

**Authors:** Trupti Shetty, Kamakshi Sishtla, Bomina Park, Matthew J. Repass, Timothy W. Corson

## Abstract

The relationship between heme metabolism and angiogenesis is poorly understood. The final synthesis of heme occurs in mitochondria, where ferrochelatase (FECH) inserts Fe^2+^ into protoporphyrin IX to produce proto-heme IX. We previously showed that FECH inhibition is antiangiogenic in human retinal microvascular endothelial cells (HRECs) and in animal models of ocular neovascularization. In the present study, we sought to understand the mechanism of how FECH and thus heme is involved in endothelial cell function. Mitochondria in endothelial cells had several defects in function after heme inhibition. FECH loss changed the shape and mass of mitochondria and led to significant oxidative stress. Oxidative phosphorylation and mitochondrial Complex IV were decreased in HRECs and in murine retina ex vivo after heme depletion. Supplementation with heme partially rescued phenotypes of FECH blockade. These findings provide an unexpected link between mitochondrial heme metabolism and angiogenesis.

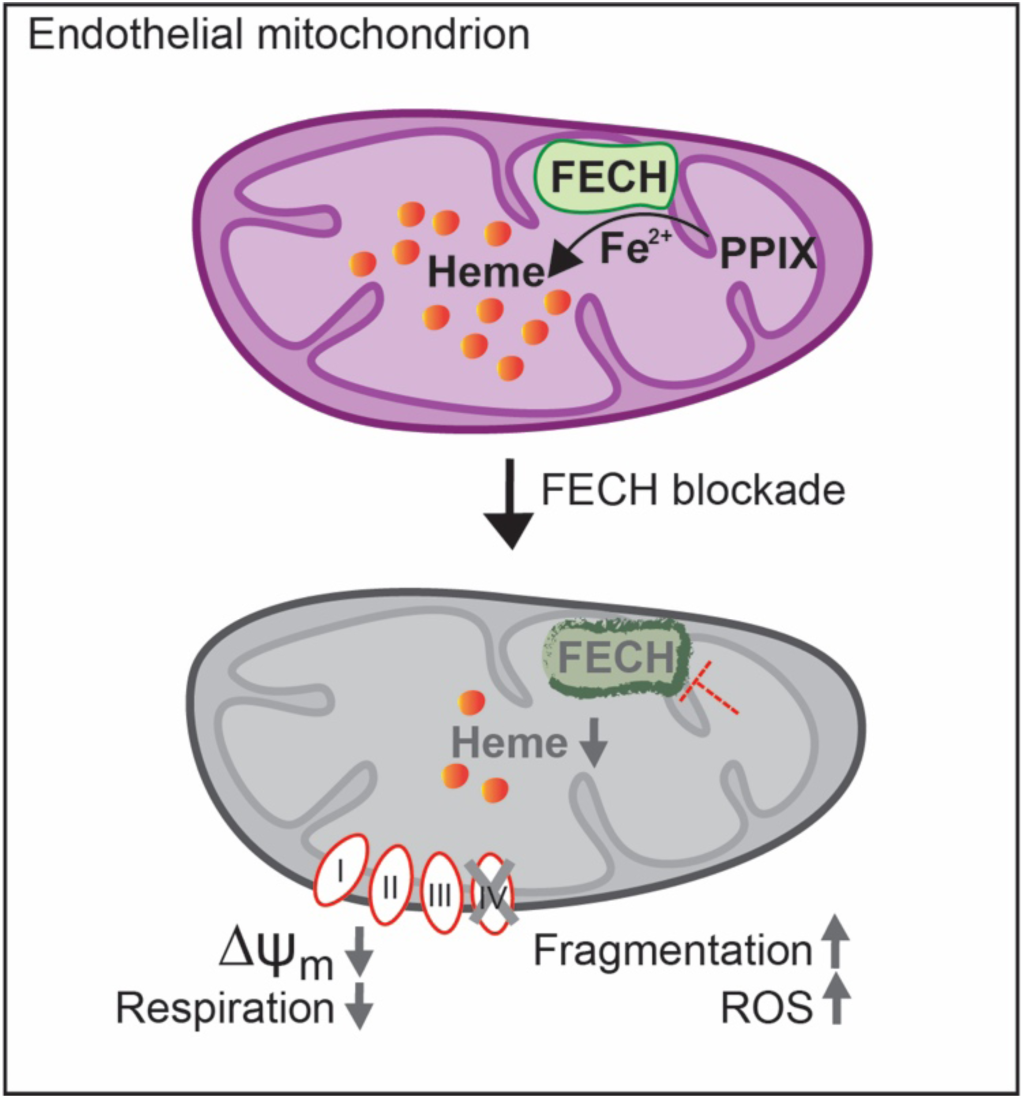

**Highlights:**

- Heme synthesis inhibition changes mitochondrial morphology in endothelial cells
- Loss of heme causes a buildup of mitochondrial ROS and depolarized membrane potential
- Endothelial cells have defective oxidative phosphorylation and glycolysis on loss of heme
- Mitochondrial damage is caused by loss of functional heme-containing Complex IV and partially restored by exogenous heme

## Introduction

An imbalance in mitochondrial metabolism has been implicated in the development of neovascular diseases catalyzed by aberrant angiogenesis. However, the role of heme synthesis in the development of mitochondrial dysfunction in neovascular diseases is unclear.

Neovascularization is a common phenomenon seen in vascular diseases like cancer, type 2 diabetes mellitus, proliferative diabetic retinopathy (PDR), wet age-related macular degeneration (AMD), and retinopathy of prematurity (ROP) (Friedman et al., 2004; Hellstrom et al., 2013; Kempen et al., 2004; Kizhakekuttu et al., 2012). Neovascularization is the disease process where new blood vessels grow from pre-existing ones and in the eye this can contribute to ocular complications like hemorrhages, retinal detachment and loss of central vision (Campochiaro, 2013). Endothelial cells (EC) are key to this process, mediating different cellular functions essential for angiogenesis like proliferation, migration and vascular permeability (Vandekeere et al., 2015).

We previously reported that heme synthesis inhibition is anti-angiogenic in retinal ECs in vitro and in animal models of ocular neovascularization. Blocking heme production in human retinal microvascular ECs (HRECs) decreased proliferation, migration, and endothelial tube formation and caused reduced protein expression of total and phosphorylated VEGF receptor 2 (Basavarajappa et al., 2017). This antiangiogenic effect was only seen in ocular ECs, while ocular non-endothelial cells were not similarly affected. Heme synthesis inhibition was associated with smaller ocular neovascular lesions in a choroidal neovascularization mouse model. However, the mechanism underlying this effect remains unknown.

The final synthesis of heme takes place in the mitochondria, when ferrochelatase (FECH) inserts ferrous ion into a precursor protoporphyrin IX (PPIX) to form protoheme (iron-protoporphyrin IX) (Dailey et al., 2017; Nilsson et al., 2009; Poulos, 2014). By directly targeting FECH, cells can be depleted of heme and heme-containing proteins, with a concomitant build-up of PPIX (Atamna et al., 2001; Vijayasarathy et al., 1999). Apart from the role of heme in oxygen transport and storage, heme acts as a prosthetic group in many hemoprotein enzymes involved in oxidative phosphorylation, plus cytochrome P450s, catalases, and nitric oxide synthase (Chiabrando et al., 2014).

In the present study, we aimed to assess the relationship between mitochondrial physiology in human ECs and heme inhibition. We hypothesized that heme inhibition would lead to defects in heme-containing enzymes of the mitochondrial electron transport chain (ETC), and thus affect mitochondrial function of ECs. We show that loss of heme altered mitochondrial morphology and dynamics, causing increased reactive oxygen species (ROS) levels and depolarized membrane potential. Our studies reveal that heme synthesis is required for EC respiration and complex IV (COX IV) function specifically, and can negatively impact glycolytic capacity of ECs. Thus, we characterize a previously unknown role of heme in cellular metabolism that facilitates EC function in angiogenesis.

## Results

### Heme inhibition caused changes to mitochondrial morphology and increased oxidative stress

To block heme production, we used a competitive inhibitor of FECH, the terminal enzyme responsible for catalyzing heme synthesis (Shi and Ferreira, 2006). Blockade of FECH with active site inhibitor *N*-methylprotoporphyrin (NMPP) causes accumulation of precursor PPIX (Figure S1A). HRECs treated with NMPP showed changes in mitochondrial fragmentation and shape (Figure 1A). NMPP treated cells had decreased form factor values, indicating reduced mitochondrial branching (Figure 1B; Figure S1B-G), owing to more highly fragmented mitochondria (Figure 1B, inset image). Mitochondria appeared less elongated and elliptical in NMPP treated HRECs as seen by lower aspect ratio values (Figure 1C). To determine mitochondrial mass, we used flow cytometry and quantified median fluorescence intensity (MFI) of NMPP treated HRECs (Doherty and Perl, 2017). NMPP treatment led to reduced MFI, indicating decreased mass in mitochondria (Figure 1D).

**Fig 1.**
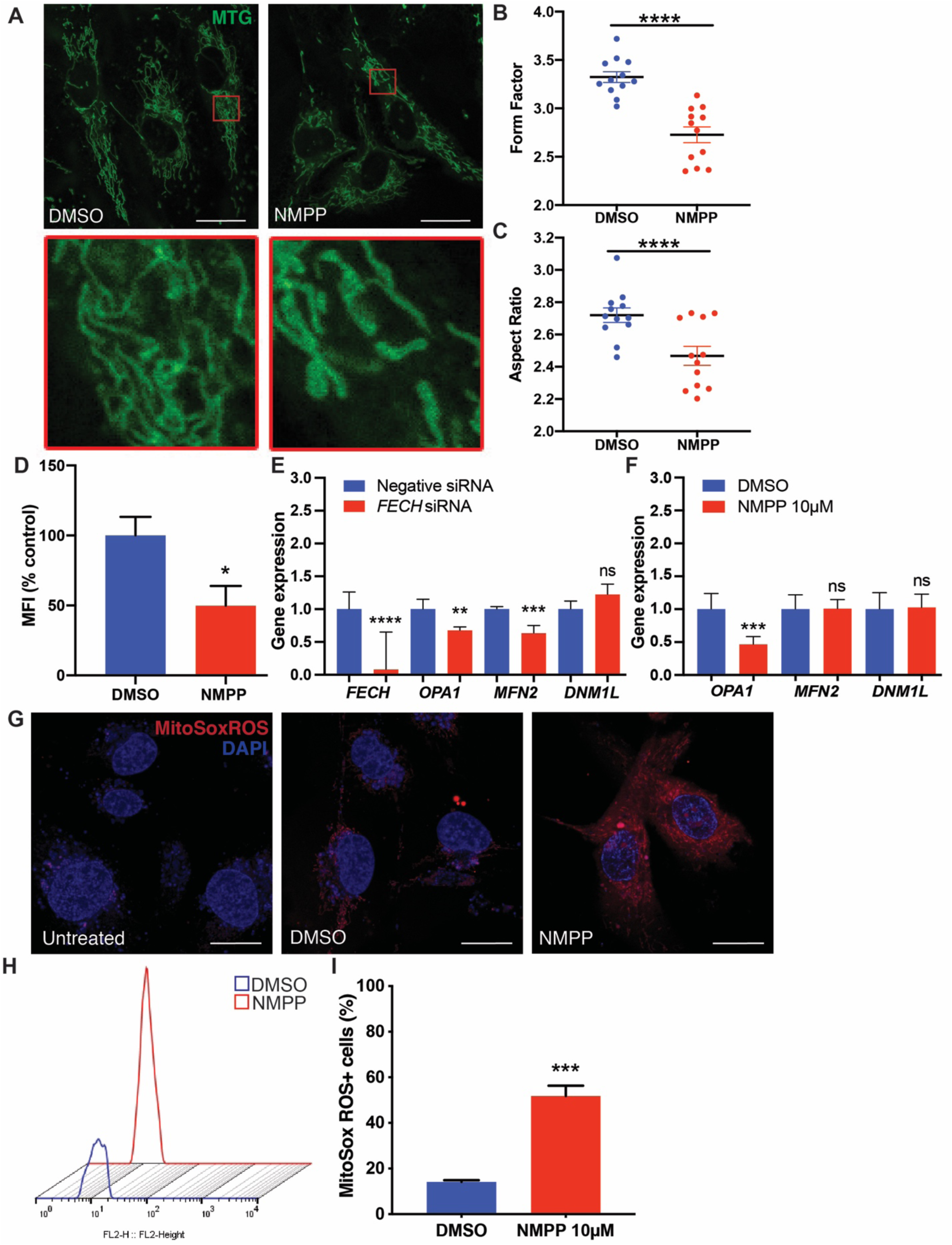
FECH blockade alters mitochondrial morphology and increases oxidative stress. **(A)** HRECs treated with DMSO or FECH inhibitor NMPP stained with MitoTracker green (MTG). Inset images indicate magnified region marked in red boxes. Form factor **(B)** and aspect ratio **(C)** as quantified using ImageJ. Individual data points indicate mean of mitochondria analyzed from each of 12 individual cells per treatment group. **(D)** Quantification of MTG fluorescence using flow cytometry and calculated Median Fluorescence intensity (MFI). qPCR analysis of mRNA expression under *FECH* knockdown **(E)** or NMPP treatment **(F). (G)** HRECs stained with mitoSox ROS in red and Hoechst staining in blue. **(H)** Representative fluorescence peaks as measured by flow cytometry followed by **(I)** quantification of cells positive for red fluorescence. Bar graphs indicate mean ± SEM, n=3. Representative results from three independent experiments. ns, non-significant, *p<0.05, **p<0.01, ***p<0.001, ****p<0.0001, two-tailed unpaired Student’s t-test. Scale bars = 20 µm.

We tested mRNA levels of key mitochondrial fusion proteins involved in mitochondrial dynamics, *MFN2* and *OPA1*, and found significantly lower levels under FECH blockade conditions (Figures 1E and 1F). Fission regulator *DNM1L* (encoding Drp1) showed no change. We measured mitochondrial specific ROS using MitoSox ROS and cells treated with NMPP showed a marked increase in ROS levels (Figure 1G). We quantified this increase using flow cytometry and found a significant increase in MitoSox ROS labeled HRECs treated with NMPP (Figures 1H and 1I). This effect of elevated ROS levels was also seen in RF/6A cells treated with NMPP (Figures S4A-B). This primate cell line has properties of chorioretinal ECs (Lou and Hu, 1987), and thus provides corroboration of the primary HREC data.

### Loss of FECH depolarized mitochondrial membrane potential

To assess mitochondrial health, we next measured membrane potential using JC-1, a polychromatic dye that on excitation forms red aggregates and green monomers depending on the energized or deenergized state of mitochondria. Both siRNA mediated knockdown of FECH and chemical inhibition using NMPP induced loss of healthy red aggregates and an increase in monomers as seen by the green fluorescence (Figures 2A and 2D). This JC-1 excitation was quantified using flow cytometry on HRECs labeled with JC-1 dye, and showed reduced red to green fluorescence ratio, indicative of mitochondrial depolarization (Figures 2B, 2C, 2E and 2F). RF/6A cells also showed depolarized membrane potential after NMPP treatment (Figures S4C-E).

**Fig. 2.**
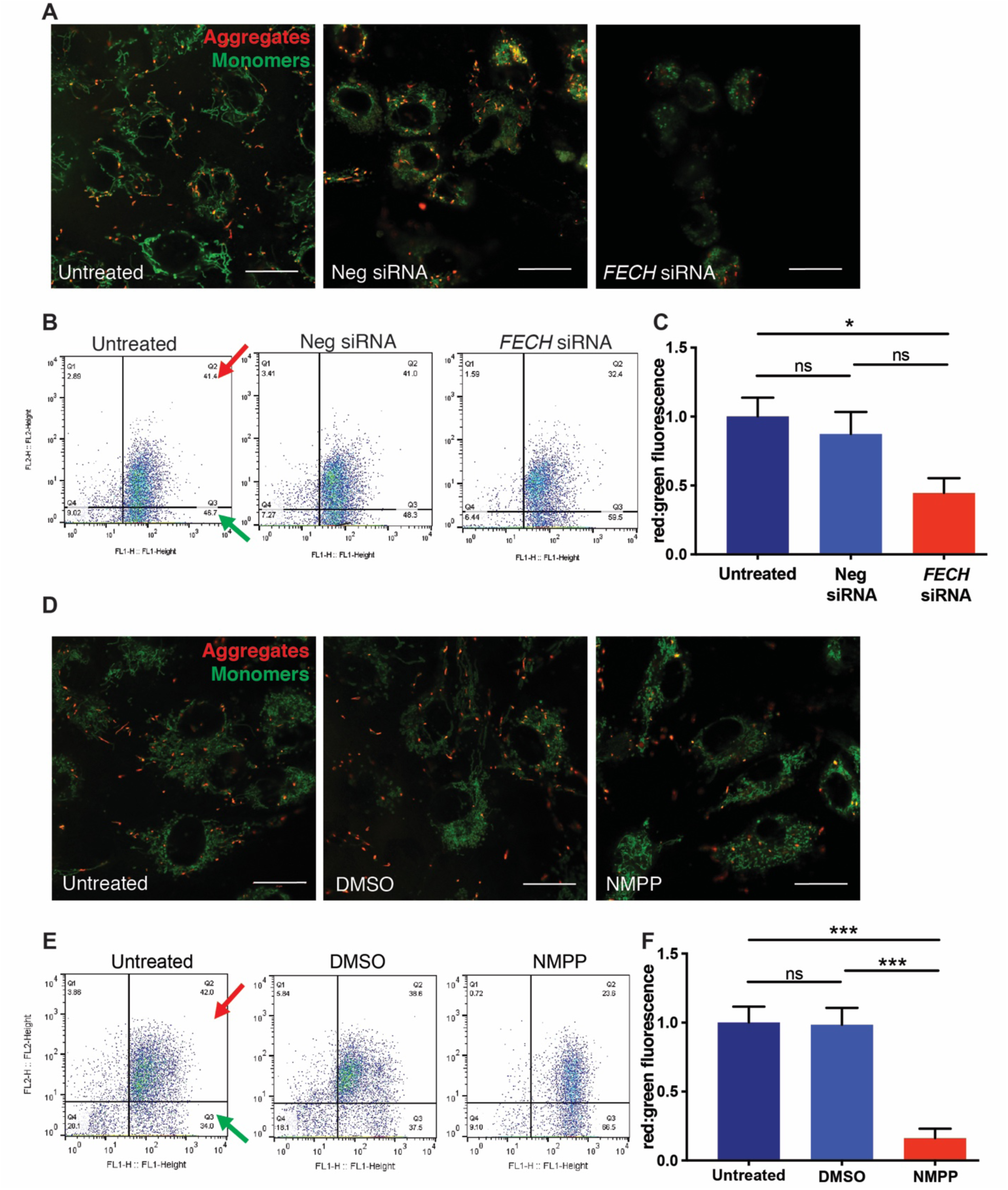
Loss of FECH reduced mitochondrial membrane potential in HRECs. HRECs stained with JC-1 dye showing green monomers and red aggregates under *FECH* knockdown condition **(A)** and NMPP treatment **(D)**. Representative dot plots of FL1 versus FL2 channel from three individual experiments, measuring red and green fluorescence using flow cytometry after *FECH* knockdown **(B)** and NMPP treatment **(E)**. Red and green arrows indicate quadrants expressing FL1-red and FL2-green fluorescent cells. **(C, F)** Quantification of red:green fluorescence from flow experiment. Bar graphs indicate mean ± SEM, n=3; ns, non-significant, *p<0.05, ***p<0.001, one-way ANOVA with Tukey’s post-hoc tests. Scale bars = 20 µm.

### Reduced COX IV expression and activity rescued by hemin

To determine where heme depletion was influencing mitochondrial function, we evaluated protein complexes of the ETC after heme inhibition, as complexes II, III and IV contain heme in their prosthetic groups (Kim et al., 2012). FECH knockdown resulted in a significant decrease of only Complex IV (Figures 3A and 3B). NMPP treated cells showed a similar decrease in COX IV, and we also found increased expression in COXV ATP synthase upon treatment with this small molecule (Figures 3A and 3C). We then examined the heme containing subunit 1 of COX IV and found a significant decrease under both knockdown and NMPP treatment (Figures 3D-G). Enzyme activity of COX IV was also reduced, and total levels of the complex were significantly reduced after FECH blockade (Figures 3H-K). To confirm if our results were dependent on heme, we sought to rescue the phenotype of COX IV reduction by external supplementation of heme to the cells. Hemin, a stable form of heme, was able to alleviate COX IV protein expression and partially rescue COX IV enzyme activity (Figures 3L-N). RF/6A cells showed a similar rescue phenotype of COX IV enzyme after heme addition (Figures S5A-D).

**Fig. 3.**
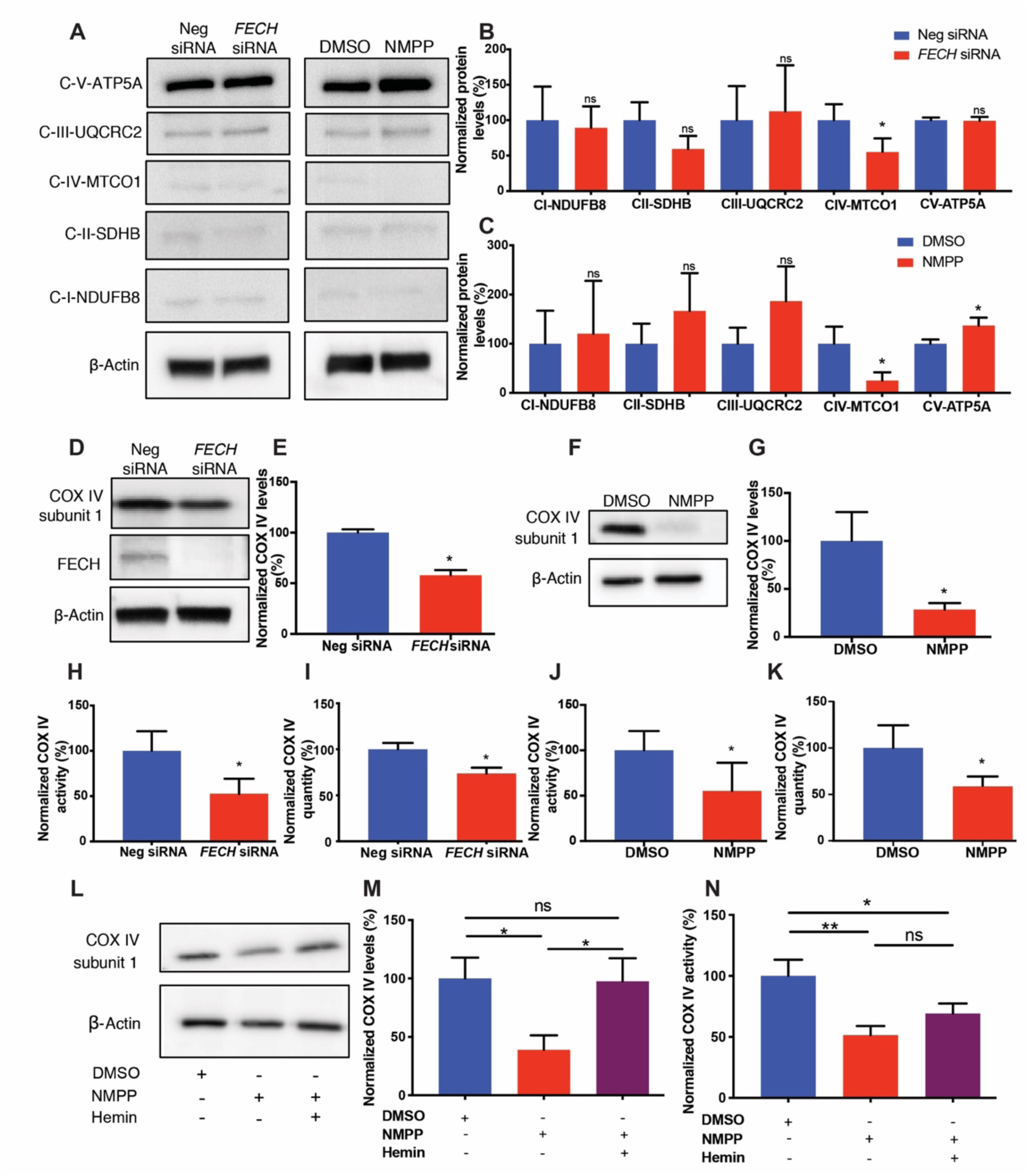
FECH inhibition caused reduced COX IV expression, rescued by hemin. **(A)** HRECs under *FECH* siRNA or NMPP treated conditions were immunoblotted for Complexes I-V as indicated. **(B, C)** Quantification of the blots was graphed as shown relative to appropriate control. **(D)** *FECH* siRNA treated HRECs were probed for COX IV subunit 1 and FECH along with housekeeping control and **(E)** quantified. Similarly, HRECs treated with NMPP were blotted for COX IV subunit 1 **(F)** with **(G)** quantification as shown. COX IV enzyme activity was measured under *FECH* knockdown condition **(H)** and NMPP treatment **(J)** and total COX IV levels were quantified by ELISA for both the conditions **(I, K). (L, M)** COX IV protein expression and quantification under defined conditions. **(N)** COX IV enzyme activity was partially rescued in NMPP treated cells exposed to hemin. Immunoblot images representative from three independent experiments. Bar graphs indicate mean ± SEM, n=3-4; ns, non-significant, *p<0.05, **p<0.01. **(B**,**C**,**E**,**G**,**H-K)** unpaired Student’s t-test **(M**,**N)** one-way ANOVA with Tukey’s post hoc tests.

### FECH blockade reduced mitochondrial function of retinal ECs

To understand whether loss of function of an important complex in the ETC resulted in defects in mitochondrial respiration, we assessed maximal OCR induced by the potential gradient uncoupler FCCP (Figures S2A, S2B). FCCP induces uninhibited flow of electrons across the ETC, causing the enzymes of the respiratory chain to use metabolic substrates at full potential and in turn, revealing the maximal cellular capacity that can meet energy demands under metabolically stressed conditions (Dranka et al., 2011). Under metabolic stress, both HRECs and RF/6A cells underwent increased glycolysis to meet energy demands (Figures S2C-F).

Based on these optimized parameters (Figure S3), we measured OCR of HRECs after FECH knockdown and observed reduced basal respiration (Figures 4A, 4B). Uncoupler-induced maximal respiration was significantly decreased, with a marked reduction also found in OCR-linked ATP production and spare respiratory capacity (Figures 4C-E). Similarly, NMPP treated cells showed a dose dependent decrease in basal (Figure 4F and 4G) and maximal respiration (Figure 4H) along with a decline in OCR-linked ATP production and spare respiratory capacity (Figures 4I and 4J). We saw a similar decrease in mitochondrial respiratory activity in RF/6A cells (Figures S5E-I).

**Fig. 4.**
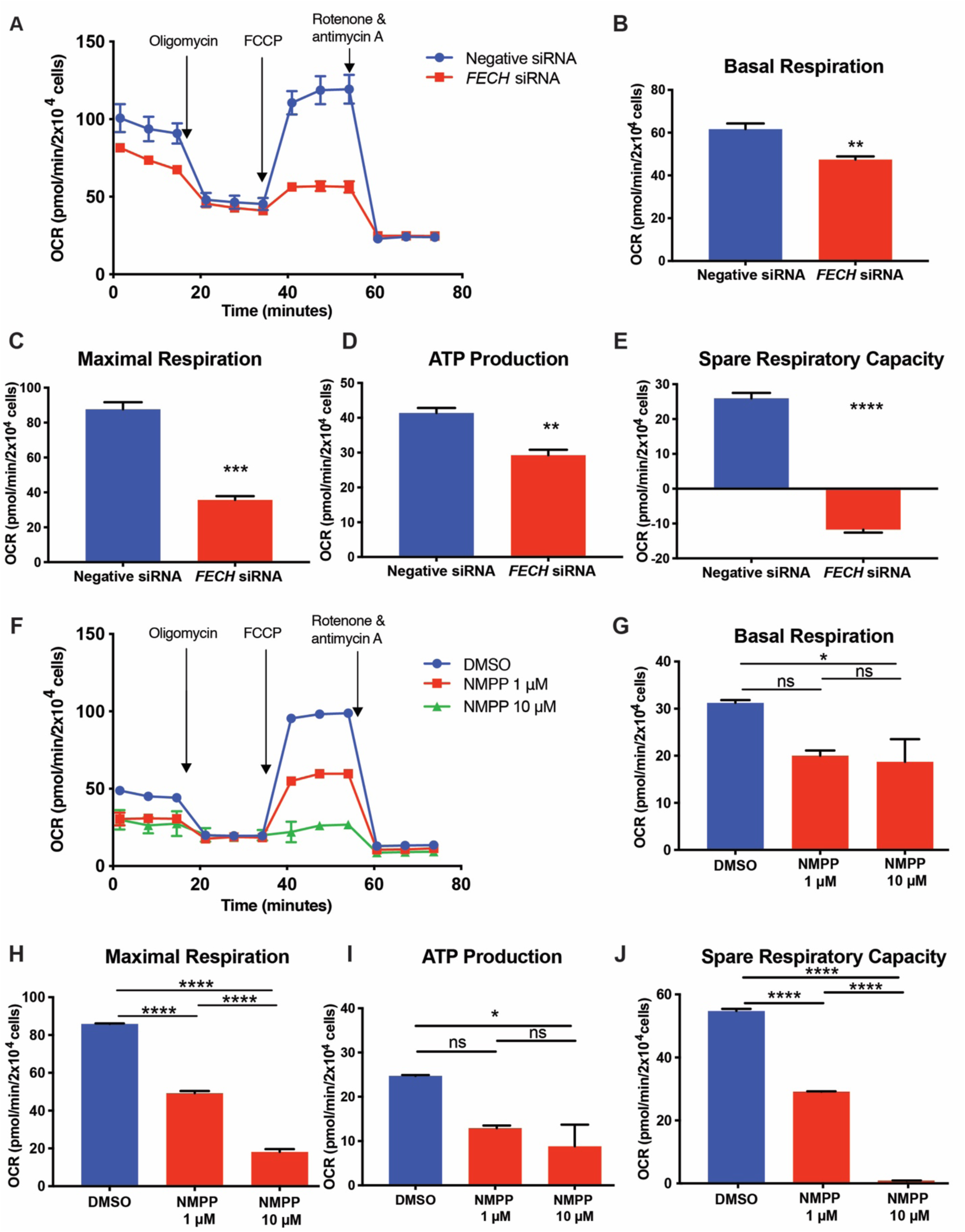
Loss of FECH reduced mitochondrial respiration. **(A, F)** OCR kinetic traces for HRECs under *FECH* knockdown or NMPP chemical inhibition. **(B, G)** Basal respiration, **(C, H)** maximal respiration, **(D, I)** OCR-linked ATP production, and **(E, J)** spare respiratory capacity were calculated based on OCR curves for the respective treatment group. **(A, F)** Representative OCR curve of three individual experiments. Bar graphs indicate mean ± SEM, n=3; *p<0.05, **p<0.01, ***p<0.001, ****p<0.0001 **(B, C, D, E)** unpaired Student’s t-test **(G, H, I, J)** one-way ANOVA with Tukey’s post hoc tests.

### Inhibition of FECH led to decreased glycolytic function

Microvascular ECs are highly glycolytic compared to other cell types (De Bock et al., 2013). Under mitochondrial stress, both HREC and RF/6A cells had increased glycolysis over mitochondrial respiration in our system (Figure S2C-F). Hence, we next investigated this key cellular energetic pathway in HRECs, by measuring changes in the pH of the extracellular medium. Cells were briefly starved in glucose deficient medium, followed by induction of glycolysis using a saturating dose of glucose. Interestingly, HRECs under siRNA mediated FECH knockdown and NMPP treatment each had decreased glycolytic capacity and glycolysis (Figures 5A-C, E-G). Both conditions depleted glycolytic reserve, as seen by a marked decrease in ECAR after 2-DG injection (Figures 5D, H).

**Fig. 5.**
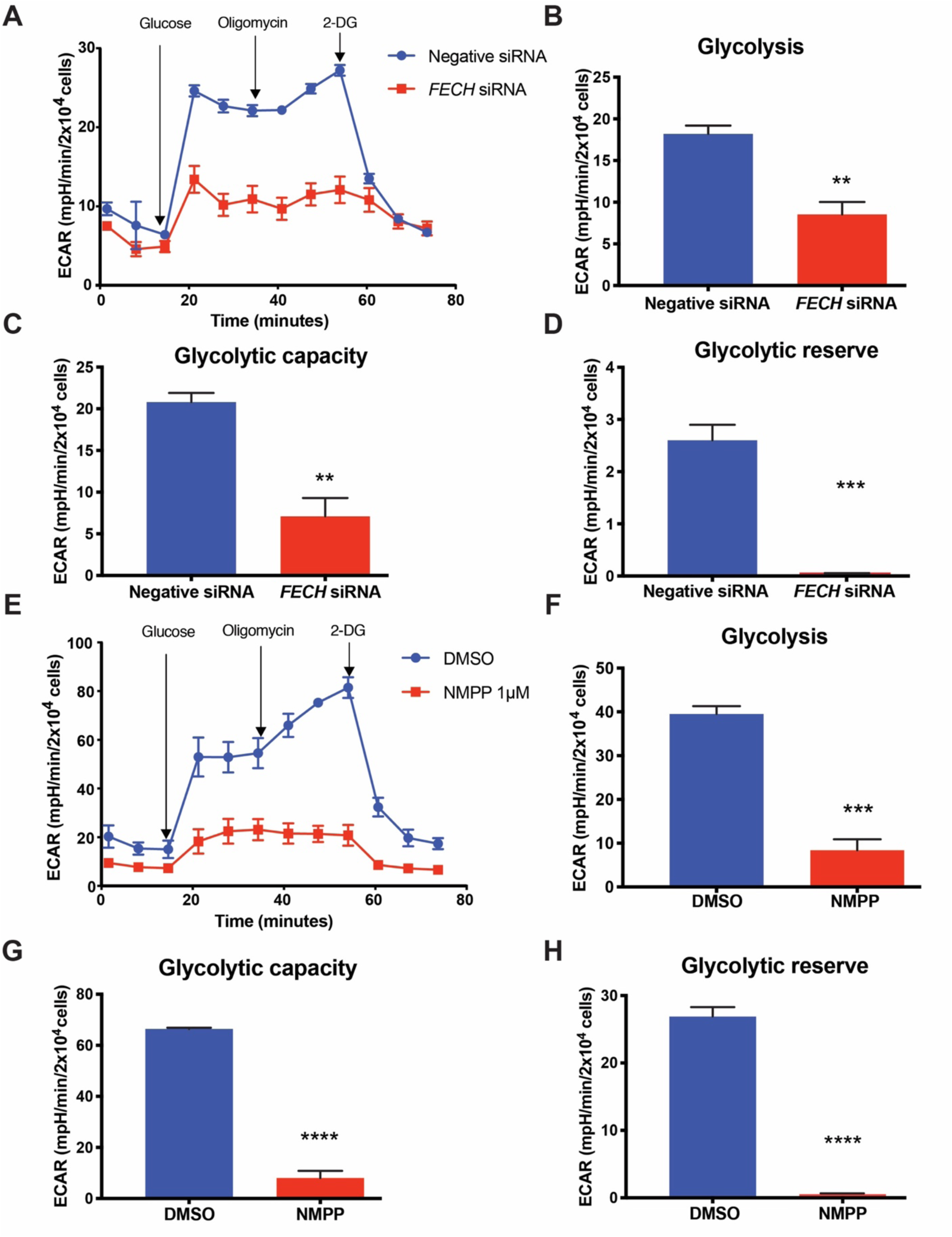
FECH inhibition caused a decrease in glycolytic function. **(A, E)** ECAR kinetic traces for HRECs under *FECH* knockdown or NMPP chemical inhibition. **(B, F)** Glycolysis, **(C, G)** glycolytic capacity, and **(D, H)** glycolytic reserve were calculated based on ECAR curves for the respective treatment group. **(A, E)** Representative ECAR curve of three individual experiments. Bar graphs indicate mean ± SEM, n=3; **p<0.01, ***p<0.001, ****p<0.0001, unpaired Student’s t-test. 2-DG, 2-deoxyglucose.

### FECH inhibition in vivo caused impaired mitochondrial energetics in retina

We further determined if the effect of NMPP had the same phenotype in vivo in an intact eye. For this, we administered NMPP intravitreally to mice and measured OCR of retina in an ex vivo assay (Figure 6A). We found that retinas treated with NMPP had a significant decrease in basal respiration and a similar decline in maximal respiration (Figures 6B and 6C). Spare respiratory capacity was severely reduced in the retinas of NMPP treated animals compared to their vehicle treated counterparts (Figure 6D). We also observed about a 30% reduction in protein expression of COX IV in the NMPP-treated retinas (Figures 6E and 6F).

**Fig. 6.**
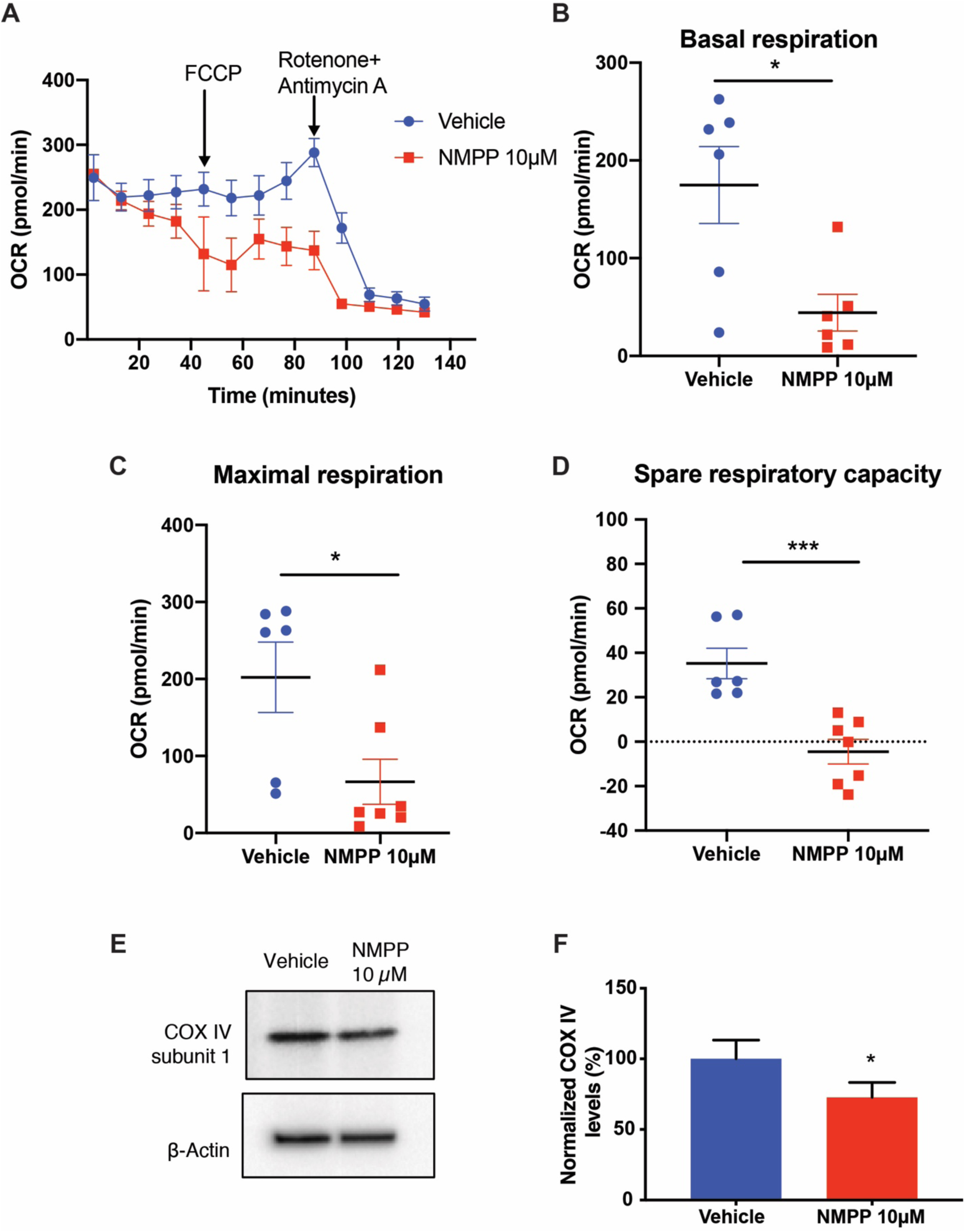
FECH inhibition in vivo decreased mitochondrial respiration in retina. **(A)** Representative OCR kinetic traces for retina from animals treated with NMPP. **(B)** Basal respiration, **(C)** maximal respiration, and **(D)** spare respiratory capacity were calculated based on OCR curves for the respective treatment groups. (**E, F**) Immunoblot showing COX IV-1 protein expression from three pooled retinal tissue lysates from NMPP treated eyes and quantification. Graphs indicate mean ± SEM for two tissue punches from each retina per treatment group, n=6-7 per treatment condition. *p<0.05, ***p<0.001, unpaired Student’s t-test.

## Discussion

Heme synthesis blockade suppresses pathological angiogenesis by poorly understood mechanisms. In the present study, we sought to investigate the effects of heme inhibition on the mitochondrial function of retinal ECs, by directly studying heme-containing complexes of the ETC and documenting mitochondrial physiology under heme depletion.

The mitochondrial electron transport chain (ETC) is a major source of ROS induced by vascular endothelial growth factor (VEGF) in hyperglycemic and hypoxic cellular environments (Cheng et al., 2011; Pearlstein et al., 2002; Wang et al., 2018). In PDR, increased mitochondrial ROS and impaired Ca^2+^ signaling can cause an increase in oxidative stress (Kowluru and Mishra, 2015; Pangare and Makino, 2012; Tang et al., 2014; Wang et al., 2018). Complex III of the ETC was recently shown to be important for umbilical vein EC respiration and thus proliferation during angiogenesis (Diebold et al., 2019). Mitochondrial dysfunction in the retinal pigment epithelium and photoreceptors has been reported for wet AMD (Barot et al., 2011; Lefevere et al., 2017), but evidence in retinal ECs is limited. Metabolic factors like succinate and adenosine, generated from the Krebs cycle and ATP metabolism respectively, are proangiogenic for hypoxia-driven neovascularization (Sapieha et al., 2010). However, the exact mechanism of how metabolites disrupt mitochondrial energetics in ischemic retinal ECs remains unclear (Grant et al., 1999; Sapieha et al., 2008).

Mitochondrial dysfunction in ECs leads to pathological angiogenesis. Heme and heme-containing enzymes play a significant role in this process, but the linkage between these phenomena has not been extensively explored. Serine synthesis deficiency induced heme depletion in ECs and caused decreased mitochondrial respiration and multiorgan angiogenic defects in animals (Vandekeere et al., 2018), while heme accumulation in ECs due to an altered heme exporter affects angiogenesis and causes endoplasmic reticulum stress (Petrillo et al., 2018). Apart from ECs, non-small cell lung carcinoma cells with elevated heme synthesis levels had increases in enzyme activities of the ETC. Increased production of heme was associated with increased migratory and invasive properties of these cells and xenograft tumors in mice (Sohoni et al., 2019). Here, we demonstrated how heme depletion by blockade of the terminal synthesis enzyme FECH led to defects in COX IV and ETC disruption (Figure 7). Retinal and choroidal ECs alike have increased mitochondrial specific oxidative stress, dysfunctional mitochondrial physiology and disruption of glycolysis as a result of these defects.

**Fig. 7.**
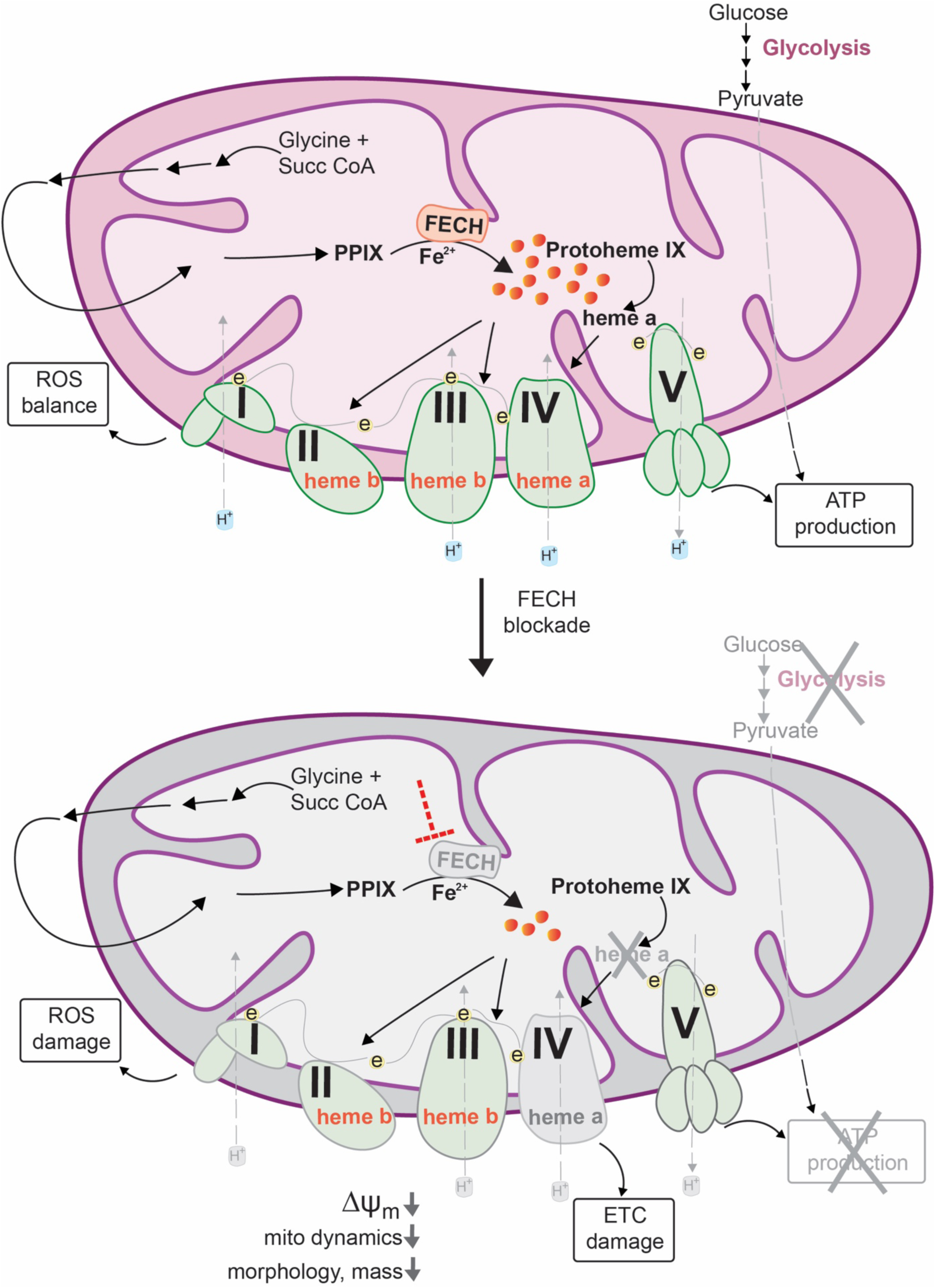
Schematic model of mitochondrial dysfunction on heme loss. Heme synthesis begins with the condensation of glycine and succinyl CoA in the mitochondrial matrix. The final step is the insertion of ferrous ion into protoporphyrin IX (PPIX) – catalyzed by ferrochelatase (FECH) to produce protoheme (also known as heme b). Protoheme and its derivatives are available for different cellular enzymes, including complexes of the electron transport chain (ETC, e = electrons). Heme a is synthesized by sub-hemylation steps, and utilized by COX IV for composition of the holoenzyme. Heme synthesis blockade by inhibiting the terminal synthesis enzyme FECH leads to COX IV defects and disrupts cellular energetics. Mitochondrial dynamics is altered with reduced fusion and mass, depolarized membrane potential (ΔΨ_m_) and elevated reactive oxygen species (ROS).

Changes to mitochondrial morphology can indicate alterations in mitochondrial dynamics and cellular function. Our findings of reduced form factor and aspect ratio after FECH blockade indicate greater fragmentation and decreased length (Duraisamy et al., 2019; Picard et al., 2013). Further, reduced mitochondrial mass was evident by our flow cytometric analysis. HRECs after heme depletion also had reduced *MFN2* and *OPA1* transcript levels. Reduction of these mitochondrial fusion protein marker genes confirms a change in mitochondrial dynamics. An increase in fragmented mitochondria also leads to increased ROS levels (Jezek et al., 2018), as confirmed by our results. Moreover, disruption of fusion proteins can also lead to an increase in ROS levels (Kim et al., 2018), as also evident in our cell model. While we did not observe any change in fission protein expression, decreased *MFN2* is also consistent with the smaller, fragmented mitochondria we observed, similar to those seen in diabetic mouse coronary ECs (Makino et al., 2010).

Inner mitochondrial membrane potential (ΔΨ_m_) was considerably decreased after FECH knockdown or NMPP treatment. This finding was explained by our result of reduced COX IV expression and activity, which might have collapsed the potential gradient and caused a reduced membrane potential. Our mitochondrial energetics findings further support this result. A significant reduction in FCCP-induced maximal respiration indicates defective COX IV, which is unable to perform the role of electron transfer. Uncoupling brought on by FCCP causes the ETC to rely on COX IV to carry out oxidative phosphorylation by the free-flowing electrons (Dranka et al., 2011). Under FECH inhibition conditions, FCCP-induced maximal respiration was affected, indicating COX IV dysfunction due to heme loss.

Heme depletion by directly blocking FECH primarily affected COX IV of all the other heme containing complexes of the ETC in ECs of the retina and choroid. COX IV subunit 1 contains heme *a* and *a*_3_ (not found in complexes I-III and V) Heme *a* is made in a series of sub-synthesis hemylation steps carried out by heme *a* synthase and is essential for proper folding and stability of COX IV (Kim et al., 2012) (Figure 7). COX IV is particularly sensitive to FECH inhibition-induced heme depletion, possibly affecting the hemylation process downstream of protoheme synthesis, leading to a smaller pool of heme *a* (Atamna et al., 2001; Sinkler et al., 2017). COX10 and COX15, proteins responsible for conversion of heme *b* (synthesized by FECH) to heme *a* were among the top-scoring genes that caused ETC disruption after heme depletion in acute myeloid leukemia cells (Lin et al., 2019), further suggesting the significance of heme *a* supply as a mediator of mitochondrial function, since it is a prosthetic group specifically in COX IV holoenzyme. Furthermore, blocking the skeletal muscle specific COX 7a subunit of COX IV decreased mitochondrial ATP production in mouse hindlimb muscles. Interestingly, this was associated with reduced muscle angiogenesis and decreased capillarity (Lee et al., 2012), further supporting a link between COX IV and EC function.

Loss of functional COX IV was restored at least partially by extracellular heme supplementation. This suggests that the EC phenotype after FECH inhibition is heme dependent and not due to an off-target effect of NMPP or due to PPIX buildup/toxicity (Wyld et al., 1997). NMPP caused significant overexpression of complex V (not seen under FECH knockdown), which could be a secondary effect of damage to COX IV, causing compensatory upregulation of ATP synthase (Havlickova Karbanova et al., 2012; Rolland et al., 2013). Our studies previously reported effects of NMPP and FECH knockdown specific to ECs of retina, choroid, brain and umbilical vein, and we did not observe a similar anti-proliferative phenotype on FECH blockade in other ocular cell types of non-endothelial origin (Basavarajappa et al., 2017). However, the phenotype of FECH deficiency on ETC complexes in other cell types beyond ECs would be an interesting aspect for future exploration.

Perhaps non-endothelial cell types can meet their energy demands from the glycolysis pathway in the event of heme depletion-induced ETC dysfunction (Rafikov et al., 2015). Conversely, loss of mitochondrial respiration in ECs due to lack of functional oxidative phosphorylation did not result in compensation by the glycolysis pathway in ECs (Zielinski et al., 2016). ECs rely upon aerobic glycolysis for nearly 85% of their energy, a feature that is highly active during aberrant angiogenesis (De Bock et al., 2013). Severe heme depletion could possibly disrupt NADH/H^+^ and redox homeostasis, as already evident from ETC dysfunction. Moreover, increased ROS levels in ECs can shift their metabolic needs to accommodate cellular damage and dysfunction, in which case both cellular energetic systems are collapsed (Vandekeere et al., 2018; Warren et al., 2014; Wellen and Thompson, 2010).

We previously tested a genetic mouse model with a M98K point mutation in the *Fech* gene (*Fech*^m1Pas^) leading to deficiency in FECH activity and found reduced neovascular lesions in the eye (Basavarajappa et al., 2017; Sardar Pasha SPB, 2019). This *Fech*^m1Pas^ mouse model shows pronounced PPIX buildup and increased mitochondrial respiratory activity, specifically COX IV (Navarro et al., 2005), likely a compensatory response to constitutive heme depletion. This renders them problematic for corroborating our findings in vivo. Moreover, complete loss of function of FECH is incompatible with survival (Magness et al., 2002). For these reasons, for in vivo validation we employed here an acute model of FECH blockade using local administration of NMPP intravitreally. Nonetheless, this FECH inhibition in vivo had energetic effects in the retina similar to those seen on retinal and choroidal ECs cultured in vitro. However, an important limitation is that this technique cannot distinguish the energetic profiles of ECs from other cell types present in the retina.

In summary, our findings provide a previously unidentified link between heme synthesis, angiogenesis, and mitochondrial energetics. The role of heme synthesis in mitochondrial function and pathological angiogenesis has been overlooked. Our observations bridge this gap in knowledge by characterizing the metabolic phenotype of ECs under heme deficiency. Loss of heme provokes prominent anti-angiogenic effects that might be exploited therapeutically for neovascular eye disease. Our findings invite future studies to further our understanding of metabolic dysfunction in neovascularization.

## Materials and Methods

### Cell Culture

Human primary retinal microvascular endothelial cells (HRECs) and Attachment Factor were purchased from Cell Systems (Kirkland, WA, USA). HRECs were grown in endothelial growth medium (EGM-2) and used between passages 4 and 8. EGM-2 was prepared by combining components of an EGM-2 “bullet kit” (Cat no. CC-4176) and endothelial basal medium (EBM, Lonza, Walkersville, MD, USA; Cat No. CC-3156). As primary cells, these were not subject to authentication. Rhesus macaque choroidal endothelial (RF/6A) cells were obtained from ATCC (Manassas, VA, USA) and grown in Eagle’s Minimum Essential Medium (EMEM, ATCC Cat No. 30-2003) supplemented with 10% fetal bovine serum (FBS, PAA Laboratories, Etobicoke, ON, Canada; Cat No. A15-751). All cells were grown at 37°C, 5% CO_2_, 100% humidity and tested for mycoplasma contamination regularly. *N*-methyl protoporphyrin (NMPP) was purchased from Frontier Scientific (Logan, Utah, USA) and prepared in DMSO. FECH siRNA (SASI_Hs01_00052189 and SASI_Hs01_00052190) was purchased from Sigma and MISSION® siRNA Universal from Sigma was used as a negative siRNA control.

### Animals

Animal studies were approved by the Indiana University School of Medicine Institutional Animal Care and Use Committee, and were consistent with the Association for Research in Vision and Ophthalmology Statement for the Use of Animals in Ophthalmic and Visual Research. C57BL/6J wild-type, healthy female mice, 8 weeks of age were purchased from Jackson Laboratories and group housed under standard conditions. NMPP at 10 µM final vitreous concentration was injected intravitreally into naïve mice under ketamine/xylazine anesthesia as described (Sulaiman et al., 2016), and 24 hrs post-injection, animals were euthanized, and retinas were immediately isolated from the animals and processed for the energetics experiment.

### Protoporphyrin IX quantification

Protoporphyrin IX (PPIX) analysis was performed at the Iron and Heme Core facility at the University of Utah. Cells were washed with PBS, pelleted, and stored frozen at - 80°C. The cells were suspended in 50 mM potassium phosphate pH 7.4 and homogenized by sonication. 200 µL extraction solvent (EtOAc:HOAc, 4:1) was slowly added to 50 µL concentration-adjusted sample and shaken. The mixture was centrifuged at 16000x*g* for 0.5 min and the supernatant was collected. About 10 µL of the supernatant solution above was injected into a Waters Acquity ultra performance liquid chromatography (UPLC) system with an Acquity UPLC BEH C18, 1.7 µm, 2.1 × 100 mm column. PPIX was detected at 404 nm excitation and 630 nm emission. Solvent A was 0.2% aqueous formic acid while Solvent B was 0.2% formic acid in methanol. The flow rate was kept at 0.40 mL per minute and the column maintained at 60°C for the total run time of 7 min. The following successive linear gradient settings for run time in minutes versus Solvent A were as follows: 0.0, 80%; 2.5, 1%; 4.5, 1%; 5.0, 80%.

### Mitochondrial morphology

Cells were plated on 35 mm coverslip bottom dishes. HRECs were stained using MitoTracker Green (Thermo Fisher, Cat no M7514) at 200 nM for 10 minutes in the dark at 37°C. Imaging was performed immediately following staining using an LSM700 confocal microscope (Zeiss, Thornwood, NY, USA) under a 63× oil immersion lens and acquired *Z*-stacked images were analyzed using ImageJ software (Trudeau et al., 2010). Briefly, individual cells were selected and particle analysis was performed to determine form factor (perimeter^2^/4*π*\*area) and aspect ratio (length of major and minor axes). Mitochondria of 12 cells per condition were analyzed and the mean per cell was considered for further statistical tests.

### Mitochondrial membrane potential assessment

Membrane potential (ΔΨ_m_) was measured with 5,5′,6,6′-tetrachloro-1,1′,3,3′-tetraethylbenzimidazolcarbo cyanine iodide (JC-1) (Santa Cruz, Santa Cruz, CA, USA) dye (Perelman et al., 2012). Cells were stained with JC-1 dye at 5 µg/mL final concentration for 10 minutes in the dark at 37°C. Cells were washed with 1× HBSS and prepared for flow cytometry (FACSCalibur, BD Biosciences, San Jose, CA) in Fluorobrite DMEM. JC-1 dye after accumulation in mitochondria forms red aggregates and green monomers that emit fluorescence at 590 nm and 510 nm respectively. For live imaging, cells grown in coverslip bottom 35mm dishes were stained and imaged using the Zeiss confocal microscope under a 63× oil immersion objective.

### Mitochondrial ROS measurement

Cells were labeled with MitoSox ROS (Thermo Fisher Scientific) dye at 5 µM final concentration for 10 minutes in the dark at 37°C using phenol red-free Fluorobrite DMEM. Cells were washed and stained with Hoechst 33342 stain for 10 minutes at room temperature. For flow cytometry, cells were labeled with dye at 1 µM final concentration, incubated in the dark at 37°C and immediately loaded for ROS fluorescence detection in the FL-2 channel using flow cytometry (Kauffman et al., 2016).

### mRNA quantification using qPCR

RNA was isolated using Trizol reagent, 1 µg RNA was used for cDNA synthesis, made using iScript cDNA synthesis kit from Bio Rad (Cat no. 1708897, Hercules, CA, USA). TaqMan probes for *MFN2* (Hs00208382_m1), *DNM1L* (Hs01552605_m1), *OPA1* (Hs01047013_m1), and *FECH* (Hs01555261_m1) were purchased from Thermo Fisher Scientific (Pittsburgh, PA, USA). qPCR was performed using Fast Advanced Master Mix, TaqMan probes, and a ViiA7 thermal cycler (Applied Biosystems). Gene expression was determined using ΔΔC_t_ method and *HPRT* (Hs02800695_m1) as housekeeping control and normalized to individual sample controls.

### Immunoblotting

Cells were lysed in RIPA buffer (Sigma-Aldrich, St. Louis, MO, USA, Cat no. R0278) and processed for protein quantification followed by SDS-PAGE. For ETC proteins, total OXPHOS Rodent WB Antibody Cocktail (Abcam, ab110413, Cambridge, MA, USA) at 1:250 dilution was used. COX IV-1 antibody was purchased from Thermo Fisher Scientific (Cat no. PA5-19471, Pittsburgh, PA, USA) and used at 1:1000 dilution. β-actin antibody was purchased from Sigma (Cat no. A5441) and used at 1:2000 dilution. FECH antibody was purchased from LifeSpan BioSciences (Seattle, WA, USA, Cat no LS C409953) and used at 1:500 dilution prepared in 5% BSA solution. Chemiluminescent reagent ECL Prime was purchased from GE Healthcare (Buckinghamshire, UK, Cat no. RPN2232).

### Complex IV activity

ELISA was performed on cell lysates treated using COX IV ELISA kit (Abcam, Cambridge, MA, USA, Cat no. ab109909) following manufacturer’s instructions. Briefly, 10 µg protein was loaded into plates coated with COX IV antibody and cytochrome *c* reduction by the immunocaptured COX IV from the sample was measured, using absorbance at 550 nm. Kinetic reads were obtained and slopes directly correlating to enzyme activity were determined. Total amount of COX IV was also assessed on the same samples.

### Hemin rescue

For hemin rescue experiments, hemin (Sigma-Aldrich, Cat no H9039) 10 µM (final well concentration) was freshly prepared in DMSO. Cells pretreated with NMPP were supplemented with hemin in EBM2 containing 0.2% FBS for 6 hrs, followed by harvesting and processing for immunoblotting or ELISA.

### FCCP titration and cell density determination

For determining optimal FCCP concentration, two ranges of FCCP were used (Figure S2A, B). Low range concentrations of 0.125, 0.25 and 0.5 µM and high range concentrations of 0.5, 1 and 2 µM were used for titration. We found 1 µM and 0.125 µM suitable for HRECs and RF/6A cells respectively. Similarly, for identification of appropriate cell seeding density to determine OCR and ECAR, cells at 5×10^3^, 2×10^4^ and 4×10^4^ cells per well were plated overnight, and photographed using an EVOS fl digital microscope, followed by Seahorse XF analyses described below.

### Mitochondrial energetics

For measuring mitochondrial function, Seahorse XFp Cell Mito Stress test kit (Cat No. 103010-100) was used (Figure S3) (Dranka et al., 2011). Mitochondrial inhibitors at final well concentrations of 1 µM oligomycin, 0.125 µM (for RF/6A cells) or 1 µM FCCP (for HRECs) and 0.5 µM rotenone/antimycin A were prepared using freshly buffered base medium. After incubating cells in a room air incubator, oxygen consumption rate (OCR) readings were determined using the XFp analyzer. The assay program was set up to measure three cycles of oligomycin followed by FCCP and a final injection of rotenone/antimycin A.

For retinal ex vivo OCR measurements, protocols were adapted as previously described (Joyal et al., 2016; Kooragayala et al., 2015). Briefly, eyes were enucleated and retina was isolated from the posterior cup. Retinal punches (1 mm diameter) were dissected from an area adjacent to optic nerve to minimize variability in retinal thickness. Retinal punches were incubated in Seahorse XF DMEM Medium pH 7.4 containing 5 mM HEPES supplemented with 12 mM glucose and 2 mM L-glutamine for 1 hour in a room air incubator at 37°C. For OCR measurement, 0.5 µM FCCP and 0.5 µM rotenone/antimycin A were injected and OCR readings were determined.

### Glycolytic function

Glycolytic parameters were measured using the Seahorse XF Glycolysis Stress test kit (Cat No. 103020-100). Cells (2×10^4^ per well) were grown overnight followed by glucose starvation in assay medium containing sodium pyruvate and glutamine for 1 hour in a room air incubator. Compounds modulating glycolysis were prepared using the same assay medium. Glucose at a final well concentration of 10 mM, oligomycin at 1µM and 2-deoxyglucose at 50 mM were loaded onto ports of the hydrated sensor cartridge followed by measurement of extracellular acidification rate (ECAR) using the XFp analyzer.

### Statistical Analysis

All Seahorse kinetic traces were analyzed using Wave 2.4 software (Agilent) and GraphPad Prism v. 8.0. FCS files for flow cytometry were analyzed using FlowJo v10. Comparisons between groups were performed with either unpaired, two-tailed Student’s t-test or one-way ANOVA with Tukey’s post-hoc tests as indicated. p values < 0.05 were considered statistically significant. Mean±SEM shown for all graphs unless indicated otherwise; n is listed in figure legends.

## Supplemental Information

Supplemental Information includes five figures S1-S5.

## Acknowledgments

This work was supported by NIH/NEI R01EY025641 (TWC). The authors thank members of the Corson laboratory for comments on the manuscript; the Indiana University School of Medicine Melvin and Bren Simon Cancer Center Angio BioCore, which is supported by NIH/NCI P30CA082790; the Iron and Heme Core facility at the University of Utah, supported in part by NIH/NIDDK U54DK110858; and the Indiana Center for Biomedical Innovation, which is supported in part by the Indiana Clinical and Translational Sciences Institute funded, in part by NIH/NCATS UL1TR002529. The content is solely the responsibility of the authors and does not necessarily represent the official views of the National Institutes of Health.

## Supplemental Figures

**Fig. S1.**
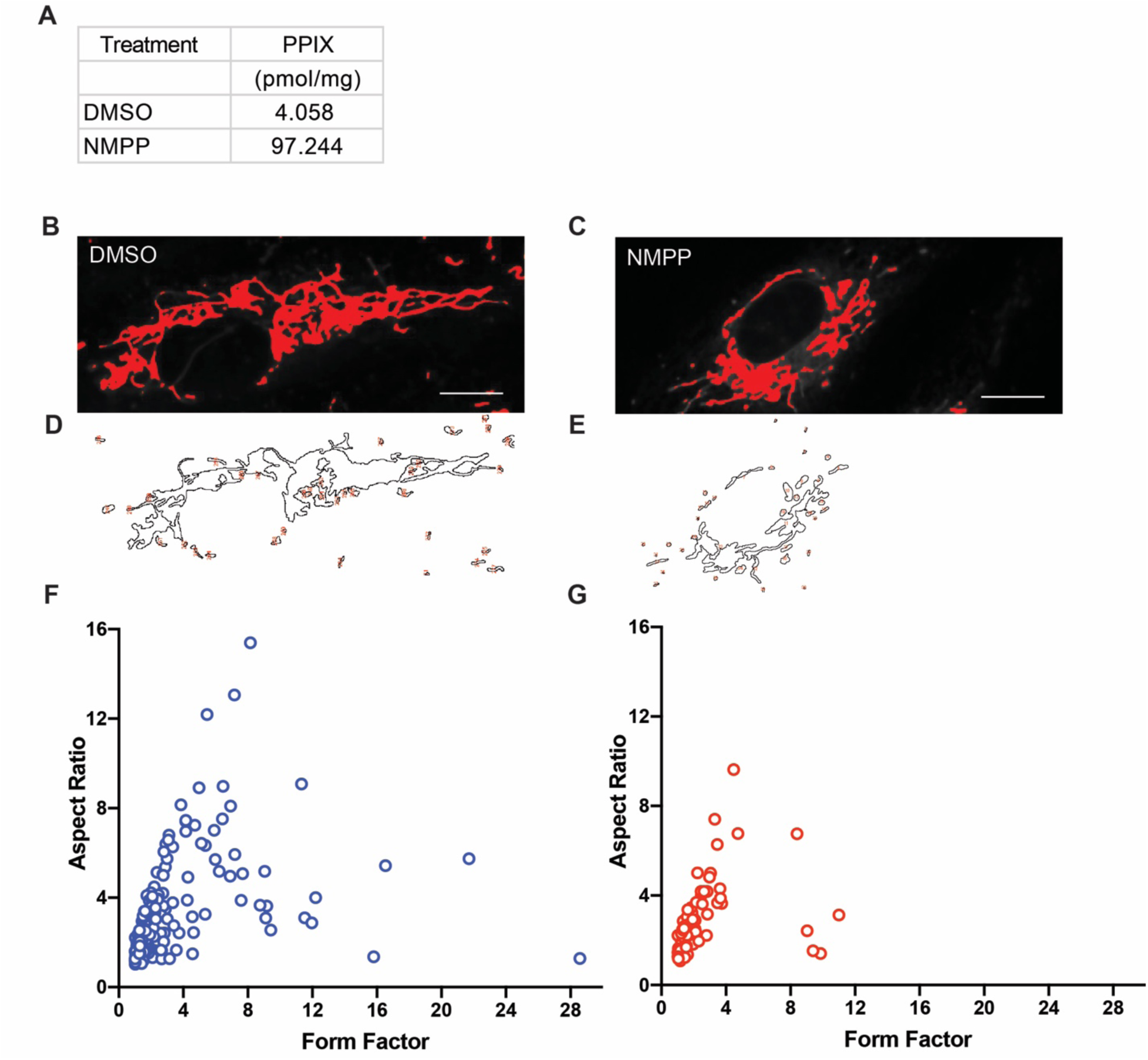
Mitochondrial morphometric analysis and PPIX levels in HRECs treated with NMPP. **(A)** PPIX levels quantified by ultra-performance liquid chromatography in HRECs treated with NMPP. Representative cells depicting particle analysis for measuring mitochondria. **(B, C)** Representative slice from Z-stack images showing threshold intensity set for DMSO and NMPP treated HRECs. **(D, E)** Representative slice from Z-stack images showing outlines of assessed mitochondria. **(F, G)** Correlation of form factor versus aspect ratio under DMSO and NMPP treatments for the representative cells shown above. Scale bars = 10 µm.

**Fig. S2.**
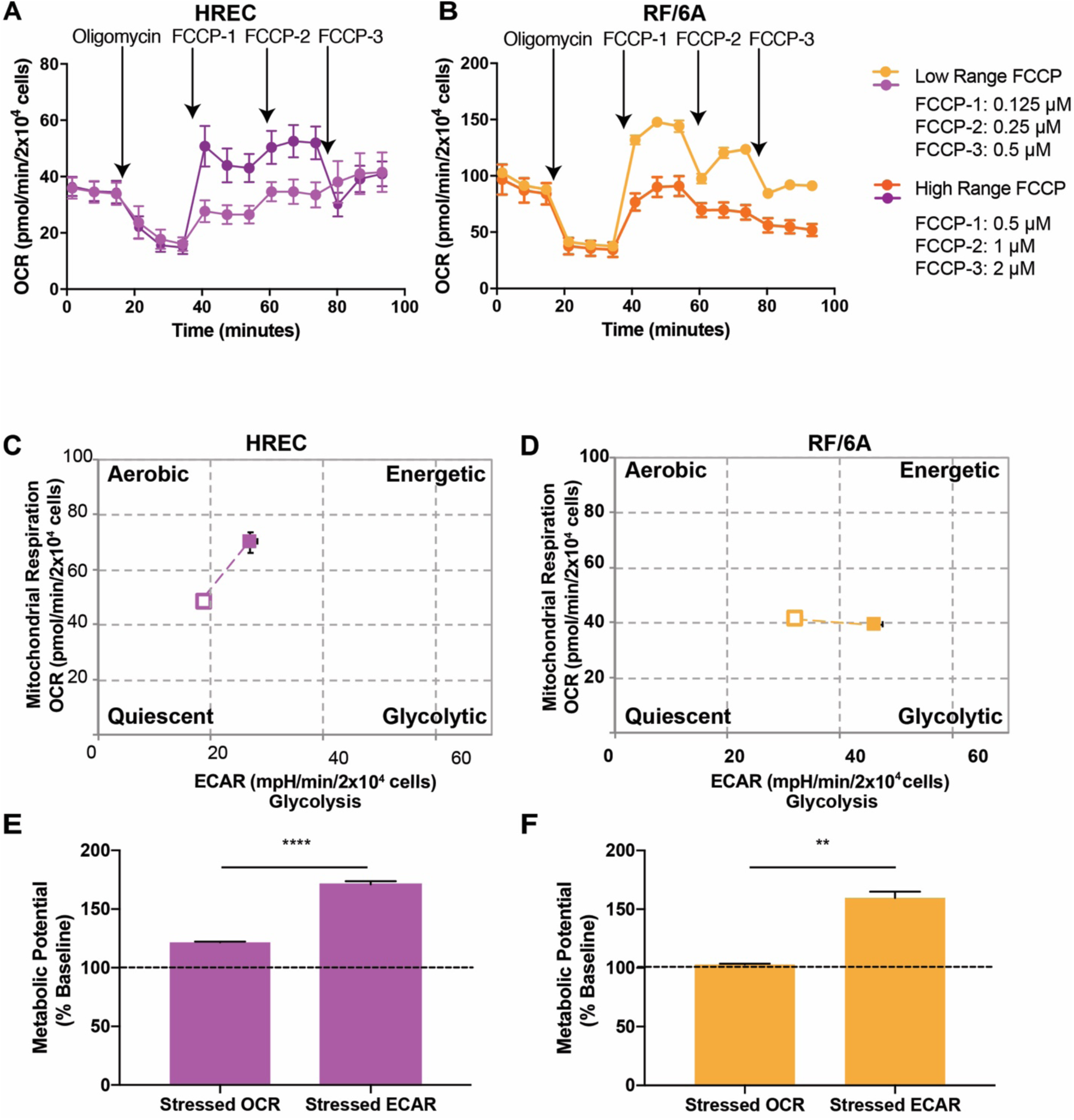
FCCP titration and cell energy phenotype of HRECs and RF/6A cells. Optimization of uncoupler FCCP concentration that produces maximal respiration in ocular endothelial cells. Oxygen consumption rate (OCR) traces of (**A**) HRECs and (**B**) RF/6A cells. 2×10^4^ cells seeded. Mean ± SEM, n=3. Cellular energetic phenotype profiles under metabolic stress of (**C**) HRECs and (**D**) RF/6A cells. Open box indicates baseline phenotype, filled box indicates stressed phenotype. Representative plots of three independent experiments. Some error bars are too small to be seen. (**E, F**) % stressed OCR and ECAR produced over 100% baseline OCR and ECAR. Mean ± SEM, n=3; **p<0.01, ****p<0.0001, two-tailed unpaired Student’s t-test.

**Fig. S3.**
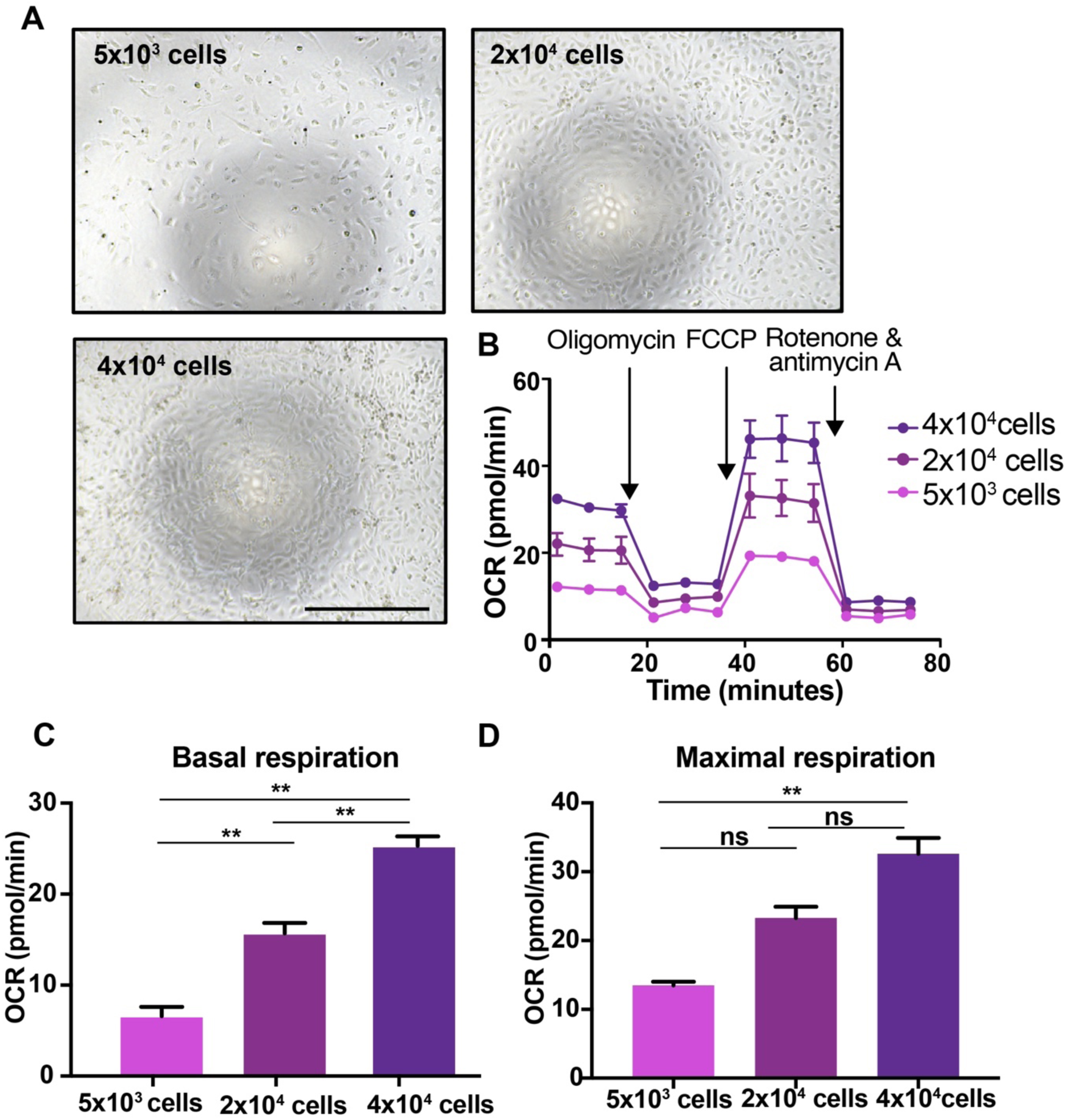
Optimizing cell seeding density. **(A)** Representative images from three independent experiments of HRECs. Scale bar = 500 µm. **(B)** Related OCR traces, **(C)** basal respiration, and **(D)** maximal respiration. Mean ± SEM, n=3; ns, not significant, **p<0.01, ANOVA with Tukey’s post-hoc tests.

**Fig. S4.**
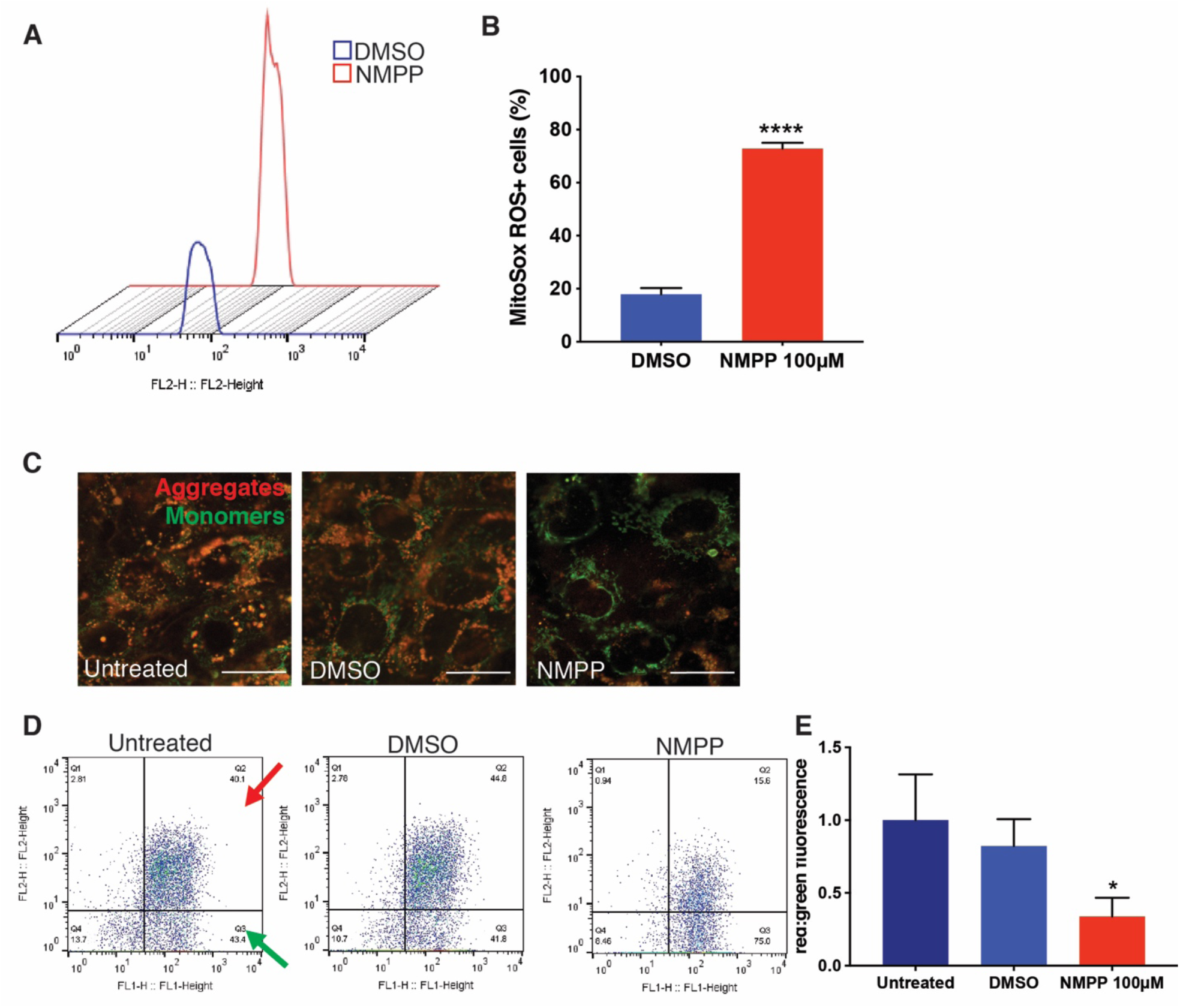
FECH blockade caused mitochondrial defects in RF/6A cells. **(A)** RF/6A cells labeled with mitoSox ROS were assessed using flow cytometry. Representative fluorescence peaks. **(B)** Quantification of cells positive for red fluorescence. ****p<0.0001 vs. DMSO, unpaired Student’s t-test**. (C)** RF/6A cells stained with JC-1 dye showing green monomers and red aggregates under NMPP treatment. **(D)** Representative dot plots of FL1 versus FL2 channel measuring red and green fluorescence using flow cytometry after NMPP treatment. Red and green arrows indicate quadrants expressing FL1-red and FL2-green fluorescent cells. **(E)** Quantification of red:green fluorescence from flow experiment. *p<0.05 vs. untreated, one-way ANOVA with Tukey’s post hoc tests. Bar graphs indicate mean ± SEM, n=3; Scale bars = 20 µm.

**Fig. S5.**
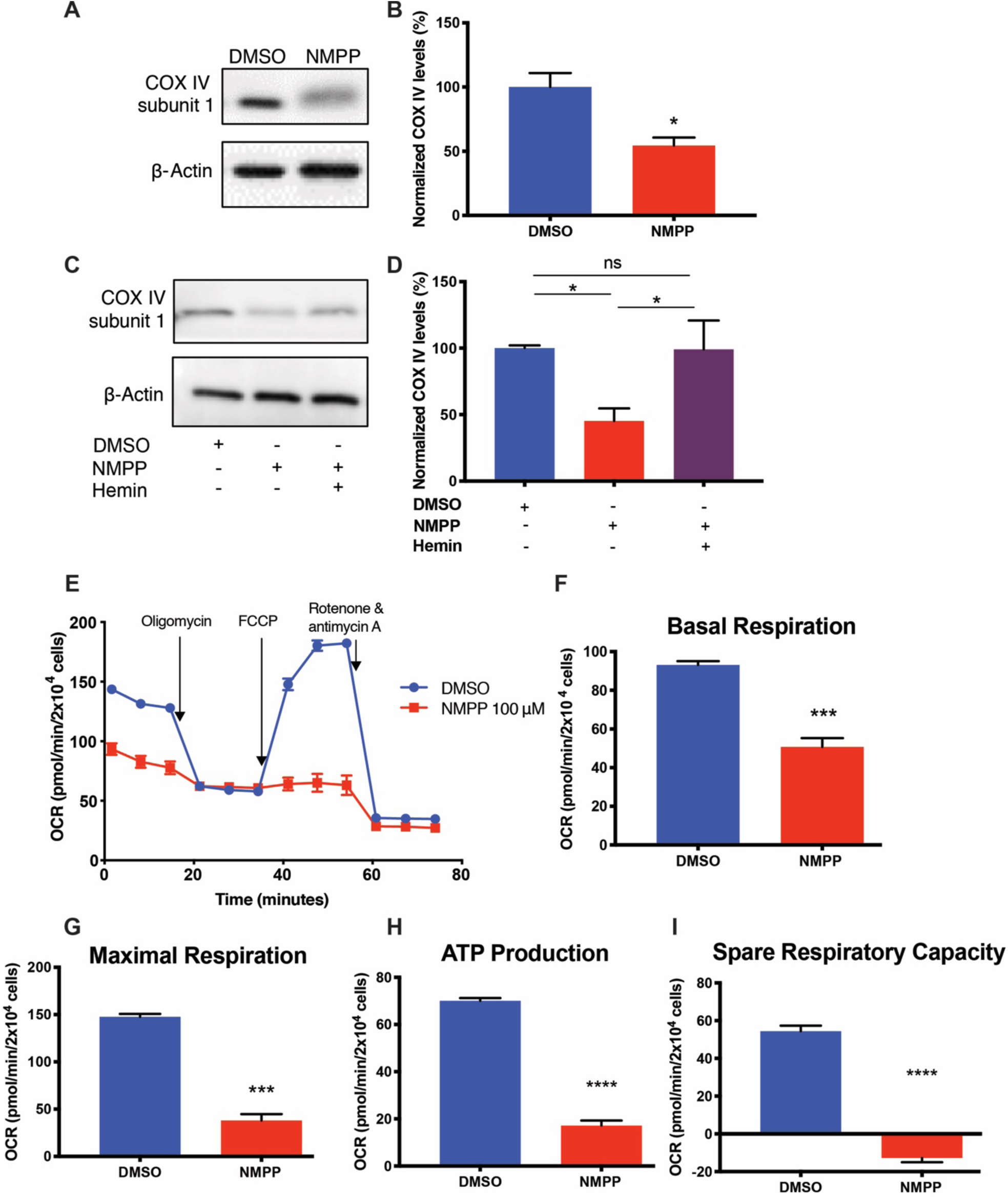
FECH inhibition decreases mitochondrial respiration and COX IV expression, which is rescued by hemin in RF/6A cells. **(A)** RF/6A cells treated with NMPP were blotted for COX IV subunit 1 with **(B)** quantification as shown. **(C, D)** RF/6A cells treated with 100 µM NMPP show rescue of COX IV protein levels after hemin supplementation. **(E)** Representative OCR kinetic traces for RF/6As under NMPP chemical inhibition. **(F)** Basal respiration, **(G)** maximal respiration, **(H)** OCR-linked ATP production, and **(I)** spare respiratory capacity were calculated based on OCR curves for the respective treatment group. Bar graphs indicate mean ± SEM, n=3; *p<0.05, ***p<0.001, ****p<0.0001, **(D)** one-way ANOVA with Tukey’s post hoc tests; (**B, F-I**) unpaired Student’s t-test.

